# On-site microbial community analysis in rivers: integrating autonomous sampling with a portable sequencing workflow

**DOI:** 10.64898/2025.12.24.695448

**Authors:** Sergio Martínez-Campos, Marc Ordeix, Maria Soria, Jose Luis Balcazar, Federico Fontana, Niccolò Rossi, Riccardo Focaia, Marzia Rossato, Massimo Delledonne, Mario Carere, Tcheremenskaia Olga, Marcheggiani Stefania, Lorenzo Proia

## Abstract

Studies focused on bacterial diversity in rivers often face significant limitations. Traditional sampling approaches frequently overlook spatial and temporal variability at the reach scale. Molecular techniques, such as metabarcoding and metagenomics, can partially address this issue by providing deeper insights. However, this approach usually involves highly specialized laboratories and high costs, making large-scale and long-term monitoring of river networks unfeasible in most cases. Thus, we developed a methodology based on remote-controlled boats, allowing the collection of integrated samples in freshwater ecosystems. Combined with a portable laboratory, this approach enabled the development of new surface water monitoring strategies. Here, we describe the operational application of this system for *in situ* monitoring of bacterioplankton communities across 8 sections of the Ter River (Catalonia, Spain). [To explore its potential, we applied both 16S rRNA gene sequencing using Nanopore technology (MinION) in the field, and shotgun metagenomics using Illumina technology in the laboratory, acknowledging the intrinsic differences between sequencing targets, platforms, and analysis pipelinesMinIONMinION sequencing enabled microbiome characterization and identification of the main bacterial taxa just 72 h after sampling, offering significantly lower costs and reduced manpower requirements. Furthermore, amplification facilitated the full characterization of bacterioplankton diversity along the river, preventing the exclusion of uncommon taxa. While shotgun metagenomics is still necessary for understanding the functional activity of these organisms, our approach provides a cost-effective framework for developing efficient follow-up sampling methodologies.

## 1. Introduction

In recent years, the increase in metabarcoding and shotgun metagenomic studies (i.e., sequencing studies of the total or partial DNA content of an environment), combined with an unprecedented drop in the price per nucleotide sequence, has enabled researchers to address fundamental questions in microbial ecology (Riesenfeld et al., 2004; Singh et al., 2009). Metabarcoding relies on the amplification and sequencing of highly conserved short DNA regions—such as the 16S rRNA gene—to provide a cost-effective and rapid approach for profiling microbial communities in complex environmental samples (Cristescu, 2014). Traditionally, this method has been performed using short-read platforms like Illumina, which offer high accuracy but require laboratory infrastructure and are limited to sequencing partial regions of the target gene (Cristescu, 2014).

More recently, portable sequencing devices such as the Oxford Nanopore MinION have enabled full-length gene sequencing directly in the field. This shift expands the potential of metabarcoding by allowing real-time, in situ microbial community analysis with a compact, field-deployable system and rapid data turnaround (Werner et al., 2022). In contrast, shotgun metagenomics sequences the entire pool of genomic DNA in a sample without prior amplification. While this untargeted approach provides broader functional insights—such as gene content, metabolic potential, and resistance profiles—it requires more sophisticated equipment, higher costs, and expert bioinformatic analysis (Quince et al., 2017). Taken together, these advances have significantly broadened the toolkit for microbial ecology. However, their implementation in routine environmental monitoring is often constrained by operational and logistical barriers. Freshwater ecosystems exemplify these challenges. Traditional monitoring methods—whether culture-based assays, targeted qPCR, or high-throughput sequencing using platforms like Illumina—typically depend on laboratory infrastructure, involve complex logistics, and result in substantial delays between sampling and data acquisition. In contrast, 16S rRNA gene metabarcoding using portable sequencing devices such as MinION provides a more accessible, cost-effective, and field-deployable alternative. By enabling rapid-time, on-site microbial profiling with minimal infrastructure, this approach has the potential to transform how microbial monitoring is conducted in freshwater environments. (Acharya et al., 2019; Hu et al., 2018; Urban et al., 2021).

Therefore, this approach may be quite relevant for monitoring and risk assessment procedures in surface waters (i.e., European Water Framework Directive) or to evaluate the microbiological quality of water bodies for bathing purposes (i.e. European Bathing Water Directive) (Wernersson et al., 2015). There are, however, some limitations to microbiological monitoring in running water ecosystems, such as the continuous river flow, which makes it necessary to increase the number of replicates, raising time-consume and relative costs, without ensuring an integrate representation of the diversity in the collected samples, or combine samples in a pool (Poretsky et al., 2014).In addition, it is necessary to filter large volumes of water to integrate adequate representation of communities, ensuring the inclusion of less abundant and potential ecologically-relevant taxa, or to sample in wide transects to achieve better representability of the ecosystem variability in terms of microbial diversity (Majaneva et al., 2018). However, despite these strategies to minimize limitations of grab-sampling based approaches some sampling sites could be discarded due to inaccessibility (Nguyen et al., 2019). Another problem is the processing of these samples which requires their handling in highly specialized laboratories, where disturbances in transport and changes in temperature can result in altered results (Acharya et al., 2020).

In this regard, portable methods, which allow for near real-time screening of water quality, could be the best option. Until now, these methods focused on the determination of bacterial concentration or the quantification of pathogenic parasites, faecal coliforms, or macroinvertebrates (Acharya et al., 2020). However, none of these methodologies ensure an extensive and integrated representation of the microbial community composition and diversity of the water body to be investigated (Acharya et al., 2020).

Instead, microbiological water quality monitoring using a portable system has been indicated as an emerging technique that could provide strong support in assessing the quality of freshwater environments (Acharya et al., 2020). This innovative approach could include the use of an Unmanned Surface Vehicle (USVs). The USVs have demonstrated to be highly effective tools for physic-chemical water quality determination, however, the development of research that tests the incorporation of new devices in the microbiological status assessment of freshwater ecosystems can be very relevant in for future environmental monitoring strategies (Katsouras et al., 2024; Mendoza-Chok et al., 2022; Warner et al., 2018). The inclusion of a filtration device with the ability to capture surface bacteria, excluding any surrounding contamination, would significantly reduce costs as well as ensure the major representativeness of spatial and temporal variability in freshwater ecosystems, and particularly in rivers (Powers et al., 2018; Preston et al., 2024). The combination of these tools with a portable laboratory that can be easily operated in the field (i.e., riverbank) to minimize the timeframe between sample collection and processing for sequencing analysis would complement this sampling approach, increasing its relevance. In addition, the use of a portable MinION sequencer with the potential to be used in the field as well, complete the novelty of this integrative approach. Portable genomic tool offer several advantages, including a comprehensive overview of microbial communities (e.g., detection of species that cannot be identified through traditional microbiological analysis), portability, and reduced costs and time of analysis (Acharya et al., 2020; Yang et al., 2021). Notably, the portability of the MinION sequencer allows for real-time analysis and use in remote locations, while requiring a low initial investment. However, it exhibits a relatively high error rate in raw sequences compared to standard NGS platforms such as Illumina (Delahaye & Nicolas, 2021; Tyler et al., 2018). Despite these limitations, the MinION platform has been widely used for monitoring water quality in well-equipped laboratory settings (Acharya et al., 2020; Hu et al., 2018; Rames & Macdonald, 2018; Urban et al., 2021). In contrast, only a few studies have used this technology in the field without reliance on complementary stationary equipment (Acharya et al., 2020; Yang et al., 2021).

We have developed and validated cost-effective microbiological monitoring of water samples in different field conditions and scenarios considering the advantages of portable sequencing devices. The procedure for using the genomic tool has been optimized and simplified so stakeholders and citizen scientists could also utilize it after appropriate training. The innovative approach included using a USV equipped with a bacteria-trapping filtration system. After that, the inclusion of a portable laboratory, allowing DNA extraction, library preparation and amplification combined with a portable sequencing device (MinION portable sequencer) with a dedicated bioinformatic analysis that allows on-site DNA barcoding to provide a comprehensive description of the water microbiome. In this context, our study aimed to describe the operational use of the portable laboratory for on-site monitoring of freshwater microbial communities and to contextualize MinION-based 16S results alongside Illumina shotgun metagenomes as complementary datasets, assessing ecological pattern concordance under a harmonized species-level framework rather than a head-to-head benchmark.

## 2. Methods

### 2.1 Methods overview

We surveyed the Ter River (Catalonia, Spain) at upstream and downstream sections using an uncrewed surface vehicle (USV) with in-line filtration, processing 0.5 L of surface water through 0.45 µm membranes; field blanks were collected and processed in parallel. DNA was extracted on site following the manufacturer’s protocol using a portable genomics setup (pipettes, mini-centrifuge, thermocycler, Qubit fluorometer, magnetic stand, and an Oxford Nanopore MinION; full inventory in Supplementary Tables S1–S2). From each extract we generated full-length 16S rRNA amplicons for MinION sequencing (flow cell and chemistry as specified), with basecalling and demultiplexing performed using ONT software unless otherwise noted, and prepared shotgun metagenomics libraries for Illumina sequencing (instrument and read length as specified) following adapter and quality trimming. Bioinformatic processing followed the MetONTIIME/ pipeline with quality filtering, chimera removal, and taxonomic assignment against a curated reference, aggregating to the species level for alpha-rarefaction and beta diversity, with higher ranks (e.g., genus/class) used only for visualization. Alpha diversity (observed richness, Shannon) and Bray–Curtis beta diversity were computed on rarefied tables; ordinations are shown as PCA. The workflow is summarized in **Figure 1**.

**Figure 1.**
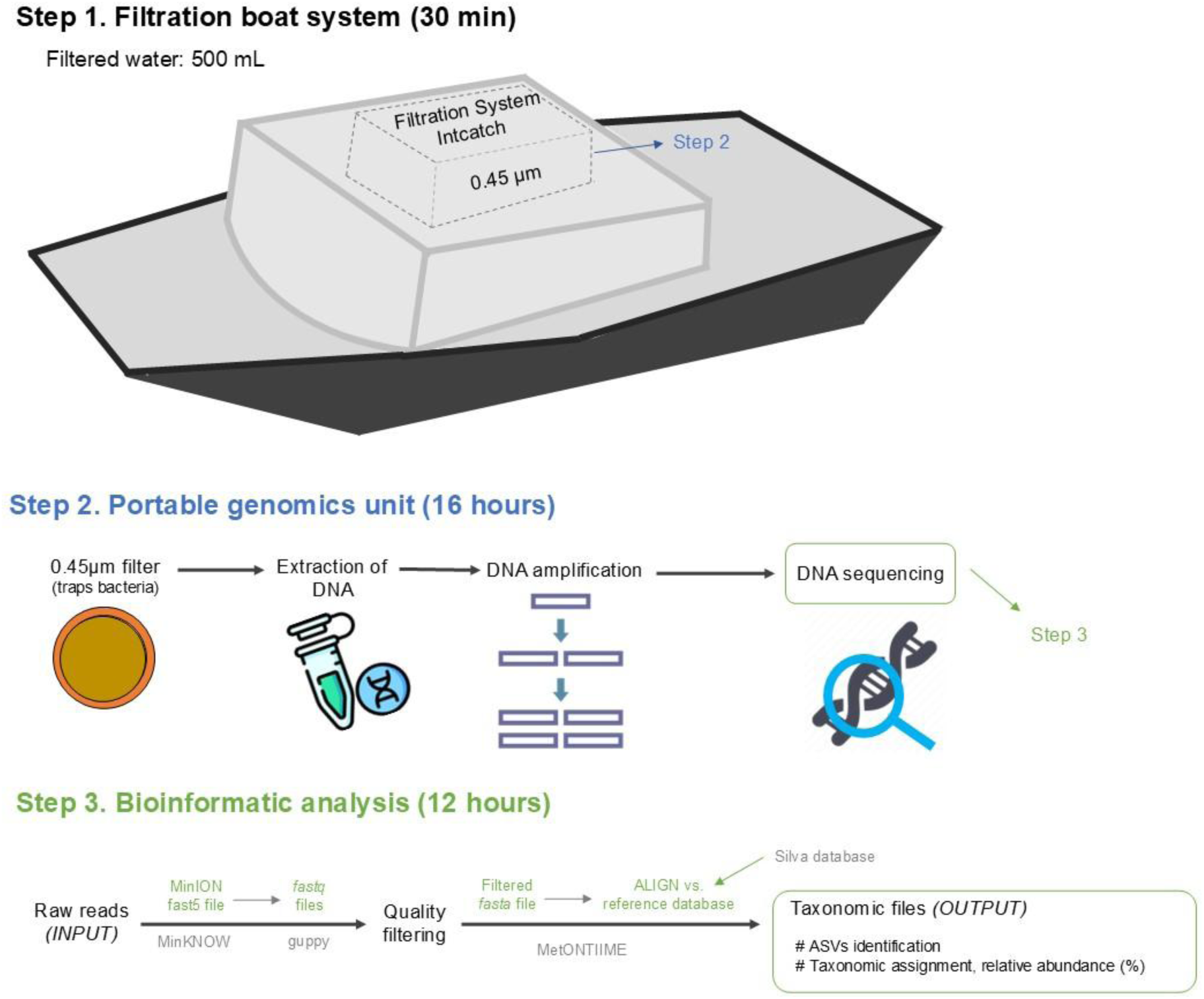
Workflow diagram of the three steps to use the portable laboratory. The boat filtration system (step 1) to improve the detection limit of the metabarcoding analysis and improve the spatial monitoring. The portable genomics unit (step 2) consists of extracting DNA from the collected 0.45 μm filter, amplifying the 16S gene, generating a DNA sequencing library and sequencing it using a MinION sequencer. Bioinformatic analysis (step 3) consists of analysing data (raw reads) using a specific pipeline and assigning the generated sequence data to taxonomic units.

### 2.2 Field sampling

Our study was conducted in May 2019 along the Ter River basin, located in the NE of Catalonia (Spain) (**Figure 2**). Ter river is divided into two sub-sections, by three main dams regulating floods and storing water reserves in the middle of its course (Jorda-Capdevila & Rodríguez-Labajos, 2015).

**Figure 2.**
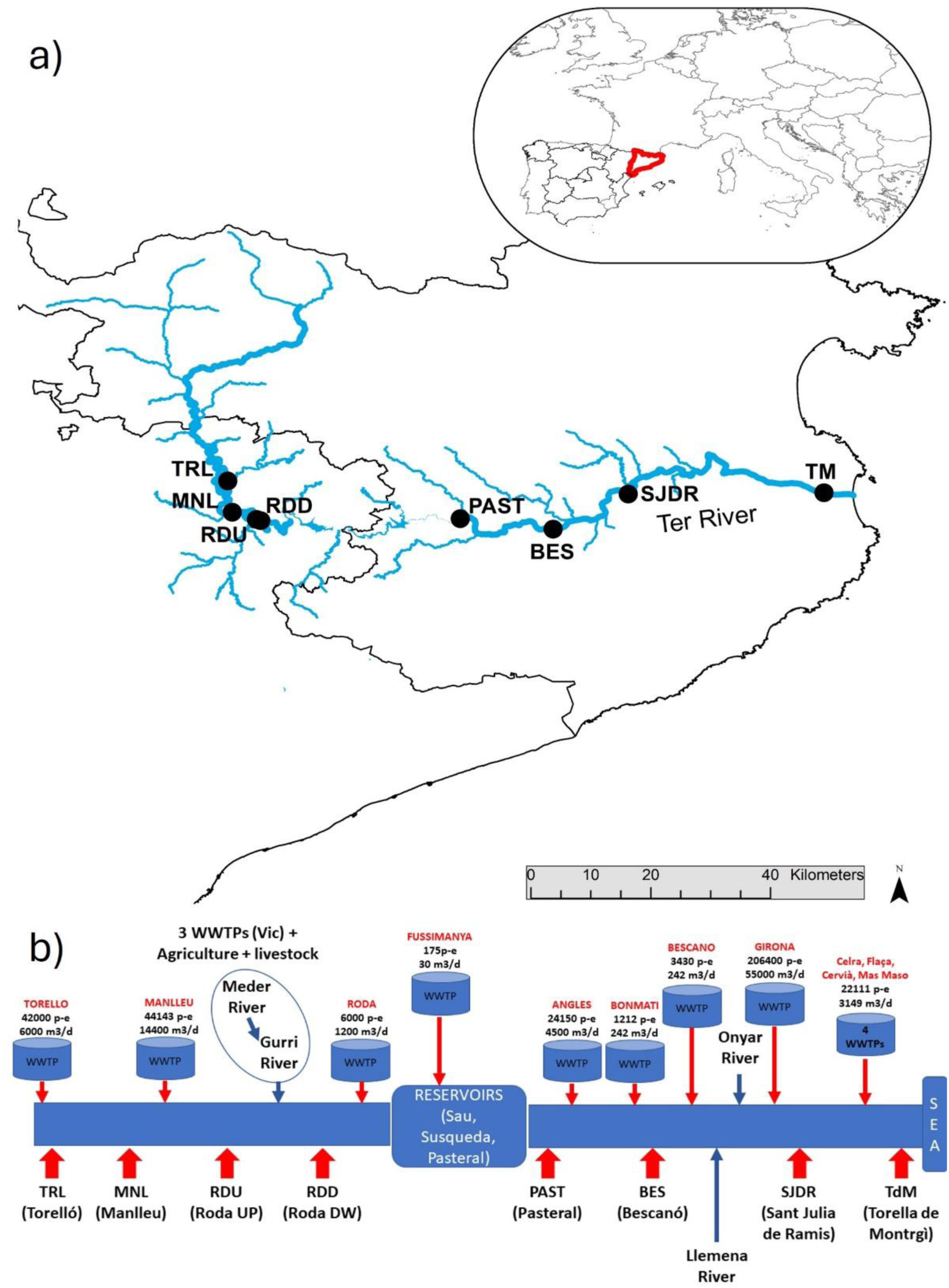
a) Sampling sites located in Ter River. b) Diagram of the main routes of pollution to the Ter River and distribution of sampling sites. Legend: TRL: Torello; MNI: Manlleu RDU: Roda Up RDD: Roda Down PAST: El Pasteral, BES: Bescano SJDR: Sant Juliá de Ramis TM: Torroella de Montgrí.

Based on this division, 4 sampling sites were selected in the upstream section, in which the dominant land use is agricultural and livestock with punctual inputs mostly from small wastewater treatment plants (Espinosa et al., 2021). These upstream sampling sites were in Torelló (TRL), Manlleu (MNL), Roda de Ter Up (RDU) and Roda de Ter Down (RDD), the latter two located upstream and downstream the confluence of between the Gurri (right tributary) and the Ter River. Subsequently, following the dams regulating floods, larger urban cores were located, which entailed the entry of a greater amount of effluent from wastewater treatment plants (Espinosa et al., 2021). In this lower section, four additional sites were selected: Pasteral (PAST) just after the last dam, Bescanó (BES) and Sant Julià de Ramis (SJDR) located up and downstream the city of Girona and Toroella de Montgrí (TDM), few kilometres before the river mouth.

### 2.3 PORTABLE WATER FILTRATION AND GENOMICS UNIT

Tools required for the portable laboratory included an USV equipped with a filtration system, a portable genomics unit (which requires access to a power supply), and an internet connection for sequencing data processing and statistical analysis. All the necessary equipment were included in **Table S1,** and fungible material for each sampling was added in **Table S2**. These items can be easily packed. The size of the filtration system allows it to be incorporated into the USV and the portable genomics unit fits into a standard suitcase (weight ∼20 kg). The timing of each step and a summary of the details can be reviewed in **Figure 1**.

#### 2.3.1 Unmanned Surface Vehicle (USVs) and Portable water Filtration Unit

The prototype of USVs employed in this project was previously described by Katsouras et al., (2024). In summary, the USVs have a total length of approximately 1 m with a width of 0.4 m; and a total weight of 14.3 kg, facilitating handling and transport. Its structure has a low profile to minimize wind resistance and can deploy in any water layer above 25 cm. The control system can be manual via remote control (at a range of 500/1000 m) or remote at a longer distance. This model featured basic measurement sensors including a conductivity meter, a dissolved oxygen probe, and a real-time pH meter, stored in flash memory and transmitted from the cloud. The electronic components were provided by GO-Systemelektronic (GO-SYS) and a BlueBox is responsible for monitoring the sensors and circuit boards (Knutz, 2018). We recorded temperature, pH, electrical conductivity, and dissolved oxygen for each transect as the set of physicochemical parameters reported in this study. Additionally, aliquots of water were collected and filtered through 0.20 μm nylon filters (Merck; Darmstadt, Germany) for the analysis of inorganic nutrients. N-NO_3_ ^-^ were analysed using ionic chromatography (Dionex IonPac AS18-Fast-4mm Column; Sunnyvale, CA, USA). P-PO_4_^3-^ and N-NH_4_^+^ were determined via spectrophotometry using a bioNova nanophotometer (IMPLEN; München, Germany).

The aquatic drone was equipped with a filtration system developed in the context of the 3.5 Task (mobile genomics laboratory) of the INTCATCH EU project. The full details of the filtration system are being published with a patent (ISS-UNIVR, PG et al.). The filtration system collected 500 mL, which was subsequently filtered to 0.45 microns.

At each sampling point, the USV was manually deployed and remotely operated. It repeatedly navigated a 100-m river transect, collecting water across the full channel width. Filtration was performed at a fixed depth of 10 cm, lasting 30 min to process 500 mL until the system automatically stopped. The procedure was repeated twice, yielding two fully saturated filters. Membrane filters were collected and then pooled prior to DNA extraction to ensure adequate yield for both the MinION amplicon and Illumina shotgun libraries; consequently, no independent per-site biological replicates were available. After each sampling, the system was sterilized and rinsed with distilled water before redeployment. Sampling proceeded from upstream to downstream to minimize cross-contamination.

#### 2.3.2 Portable genomic unit

The portable genomics unit allows the entire DNA metabarcoding procedure to be performed, from DNA extraction to sequencing and taxonomic classification of bacteria (**Figure 1**). This tool consists of three micropipettes (P1000, P200 and P20, Eppendorf), a mini-microcentrifuge (Prism Mini Centrifuge, Labnet, Cat. No. C1801230V-EU), a portable thermal cycler (MinION Systems, Cat. No. M4001), a fluorometer (Qubit 4, Cat. No. Q33240, Thermo Fisher Scientific), a vortex (Scientific industries Inc., Cat No. SI0256 230V) with Qiagen adapter (Cat No. 13000-V1-5), a magnet for bead-based purifications, the nanopore sequencer (MinION, ONT) and an ASUS laptop (Intel® i7 CPU, 2.6 GHz, 4 cores with hyper threading, 16GB RAM and 512GB SSD). All The equipment was transported in a single protective case (55×45×20 cm, Peli, Cat No. 1615) and reagents that required stored at 4 °C or –20 °C were transported in a foam box containing ice packs. Furthermore, the experiment required the use of a small generator (minimal potency of 2000W)

Once the equipment is assembled, the following 3 steps have to be performed: (1) extraction of bacterial DNA from from the 0.45 uM filter was performed using DNeasy® PowerWater® Kit (Qiagen Cat No. 14900-50-NF); (2) 16S rRNA gene amplification and library preparation for ONT sequencing was performed from the extracted DNA using using the 16S amplicon barcoding kit (SQK-RAB204) following the manufacturer’s instructions and (3) sequencing was performed on a R9.4.1 flow-cell® (FLO-MIN106, Oxford Nanopore Technologies, London, UK) using a MinION® Nanopore sequencer (Oxford Nanopore Technologies, London, UK) for ∼7h. The software MinKNOW was used to control the sequencing operation and obtain the fast5 files.

In the case of shotgun metagenomics sequencing, the DNA extracted during the first step was stored at −20 ᵒC. Subsequently, the quality and quantity of DNA samples were assessed using agarose gel electrophoresis and a VICTOR Nivo multimode plate reader (PerkinElmer; Waltham, MA), respectively. These samples were then processed using Illumina TruSeq DNA PCR-free libraries and sequenced on the NovaSeq 6000 platform (Illumina Inc., San Diego, CA) with 150-bp paired-end reads.

Sequences used in this study were submitted to the National Center for Biotechnology Information (http://www.ncbi.nlm.nih.gov/) under the Bioproject accession number: PRJNA1218628.

#### 2.3.4 Bioinformatic analysis

To harmonize taxonomic resolution across approaches, 16S rRNA profiling analysis was performed.

In the case of Nanopore sequencing, 16S rRNA profiling was performed using the MetONTIIME pipeline (Matoute et al., 2024). In the first step, the Guppy (V4.0.11) software performed the base-calling and converted the data to FASTQ format. After that, the use of MetONTIIME allowed the whole bioinformatics process to be developed in a single step up to the taxonomic classification. For filtering, the –minQual option was used to retain reads with a minimum per PHRED quality value of 12, --MinReadLenght and -maxReadLenght were used to keep sequences size between 1400 and 1500 bp. Then, the taxonomical classification was performed using VSEARCH (Rognes et al., 2016) with Silva v132 database (Quast et al., 2013). To equalize depth across samples, each library was randomly downsampled to 100,000 reads prior to diversity analyses (seed=42 without replacement). MetONTIIME (≥v2.0.0) was executed under Nextflow to improve process-level reproducibility. Exact command lines, software versions, and SILVA reference filenames with MD5 checksums are given below and mirrored in Supplementary Protocol S1.For shotgun metagenomic sequencing, the read files were converted to FASTQ format using the bcl2fastq package. The raw sequences were then filtered using FASTX-Toolkit, requiring at leat 90% of bases with PHRED ≥ 20 per read The resulting filtered reads were assigned to obtain the compositional 16S rRNA sequences using METAXA2 (Bengtsson-Palme et al., 2015) for taxonomic classification and data normalization.From the taxonomic classifications obtained by both methods, taxonomic profiles were collapsed to the genus level for comparability, and the relative abundance of the lowest taxonomic classifications was used to calculate the rarefaction curves, α-diversity and β-diversity. The rarefaction curves was computed at the species level from per-sample count tables after filtering and curated taxonomic assignment. α-diversity was calculated from the Shannon-Weaver index (Shannon, 1948) and the β-diversity was represented by a PCA after calculating the Bray–Curtis dissimilarity matrix (Bray & Curtis, 1957) using the Vegan package and ggplot2 (Dixon, 2003; Wickham, 2016) in R.

## 3. Results

### 3.1 Field implementation: logistical and economical aspects

This study demonstrated the end-to-end feasibility of the workflow under field conditions. Beyond sequencing outputs, the campaign revealed actionable logistical, operational, and economic considerations for future deployments (**Figure S1** for check images of the process). We deployed a single uncrewed surface vehicle (USV) with in-line filtration and completed eight sequential collections across two river transects, recovering the boat between sites to swap filter holders and log metadata. Because the workflow was designed to be self-contained, neither running water nor a clean room was required; we operated from a folding table under a shade canopy using disposable bench pads, bleach/70% ethanol wipes, and single-use consumables. The portable genomics lab fit into one protective case (∼20 kg; 55 × 45 × 20 cm) and was flown as standard luggage.

Field operations were handled by a two-person team (USV operator and bench operator). At each site, the USV filtered ∼500 mL over ∼30 min at a controlled flow before recovering the filter set for DNA extraction; this volume/time combination was optimized within INTCATCH. Sequencing runs on a MinION typically lasted ∼7 h, enabling preliminary outputs within ∼12–24 h when combined with on-site library preparation and laptop-based analysis. Mobilization (equipment checkout, reagent pre-aliquoting, packing) and pack-down/decontamination required ∼1.5 h each; the eight-site campaign totaled ∼30 person-hours.

Contamination control was addressed through field blanks, pre-aliquoted reagents, and physical separation of clean and dirty zones. Two waste streams were generated across all sites: non-hazardous solid waste (filters, tubes, tips, gloves, wipes) totaling ∼10–12 L (0.5–0.8 kg), and liquid waste that was negligible (≤250 mL of disinfectant/rinse combined); all waste was returned to the laboratory for disposal.

The consumables cost of metabarcoding a single water sample with the portable genomics lab was ∼€900 per sample, with the majority (∼90%) attributable to DNA amplification and sequencing; multiplexing 12 samples on a single flow cell reduced the cost to ∼€93 per sample. **Table 1** summarizes reagents and per-sample consumables for both scenarios (consumables only **Table 1**; the rest of the material can be checked in **Table S1** and **Table S2**).

**Table 1.** Breakdown of consumables cost per sample for the workflow (filtration/extraction and 16S–MinION amplification/sequencing), listing supplier, unit cost (€), and cost per sample for a single-sample run (n=1) and for 12 samples processed in parallel (n=12). In the n=12 scenario, a single R9.4 flow cell and 16S Rapid Amplicon Barcoding Kit are shared across 12 samples

| Stage | Item | Supplier | Unit cost (€) | Cost per sample (1 sample run (€)) | Cost per sample (12 samples in parallel (€)) |
| --- | --- | --- | --- | --- | --- |
| Filtration & DNA extraction | Filter 180 µm Ø 47 mm | Merck Millipore | 1.28 | 1.28 | 1.28 |
|  | Filter 20 µm Ø 25 mm | Merck Millipore | 1.07 | 1.07 | 1.07 |
|  | Filter 5 µm Ø 25 mm | Merck Millipore | 1.07 | 1.07 | 1.07 |
|  | Filter 0.45 µm Ø 25 mm | Merck Millipore | 1.17 | 1.17 | 1.17 |
|  | Qubit dsDNA BR Assay Kit | Thermo Fisher | 1 | 1 | 1 |
|  | DNeasy PowerWater Kit | Qiagen | 10.28 | 10.28 | 10.28 |
| DNA amplification & sequencing | 16S Rapid Amplicon Barcoding Kit | Oxford Nanopore | 92 | 92 | 7.67 |
|  | Agencourt AMPure XP | Beckman | 0.5 | 0.5 | 0.5 |
|  | LongAmp Taq | New England Biolabs | 2.08 | 2.08 | 2.08 |
|  | SpotON Flow Cell R9.4 | Oxford Nanopore | 800 | 800 | 66.67 |
| TOTAL CONSUMABLE COSTS (€/sample) |  |  |  | 910.45 | 92.79 |

### 3.2 Environmental parameters

Conductivity showed a gradient among locations. The maximum was recorded at TM (664 µS cm⁻¹), followed by SJDR (485 µS cm⁻¹) and RDD (407 µS cm⁻¹), with intermediate values at BES (392 µS cm⁻¹) and the MNL–RDU–PAST block (360 µS cm⁻¹). The minimum corresponded to TRL (312 µS cm⁻¹). Comparatively, TM approximately doubled TRL, evidencing a marked increase toward the lower reach of the system.

Temperature was highest at TM (26.3 °C) and MNL (26.2 °C), with similarly high values at RDU (24.9 °C) and RDD (24.7 °C). In contrast, PAST showed the lowest temperature (12.2 °C), followed by BES (14.1 °C) and SJDR (18.1 °C). The overall range among codes reached 14.1 °C. pH remained generally alkaline, with maxima at MNL (8.89) and RDU (8.85), high values at TRL (8.77) and RDD (8.76), and lower values at TM (8.57), BES (8.39), and SJDR (8.33); PAST showed the minimum (8.12).

For dissolved oxygen, the highest concentration was observed at BES (11.6 mg L⁻¹), followed by SJDR (10.7 mg L⁻¹) and MNL (9.9 mg L⁻¹), while PAST recorded the minimum (8.3 mg L⁻¹). Oxygen saturation reached its maxima at TM (125.4%) and MNL (123.2%); values at SJR (113.2%) and BES (113.1%) were also high, whereas PAST presented the lowest saturation (77.8%).

Regarding nutrients, NH₄⁺-N was generally low, with RDD showing the highest value (0.18 mg L⁻¹), followed by RDU (0.11 mg L⁻¹); the remaining codes ranged 0.07–0.08 mg L⁻¹. For NO₃⁻-N, maxima were found at PAST (2.97 mg L⁻¹), BES (2.86 mg L⁻¹), and SJDR (2.72 mg L⁻¹), with intermediate values at TM (2.34 mg L⁻¹) and minima at RDU (0.59 mg L⁻¹) and MNL (0.74 mg L⁻¹). PO₄³⁻-P showed a distinct pattern, with maxima at RDU (216 µg L⁻¹) and RDD (198 µg L⁻¹), above TM (96 µg L⁻¹) and TRL (76 µg L⁻¹), and minima at BES (∼40 µg L⁻¹) and PAST (∼39 µg L⁻¹). A summary of the physicochemical and nutrients parameters is provided in **Table 2**.

**Table 2.**
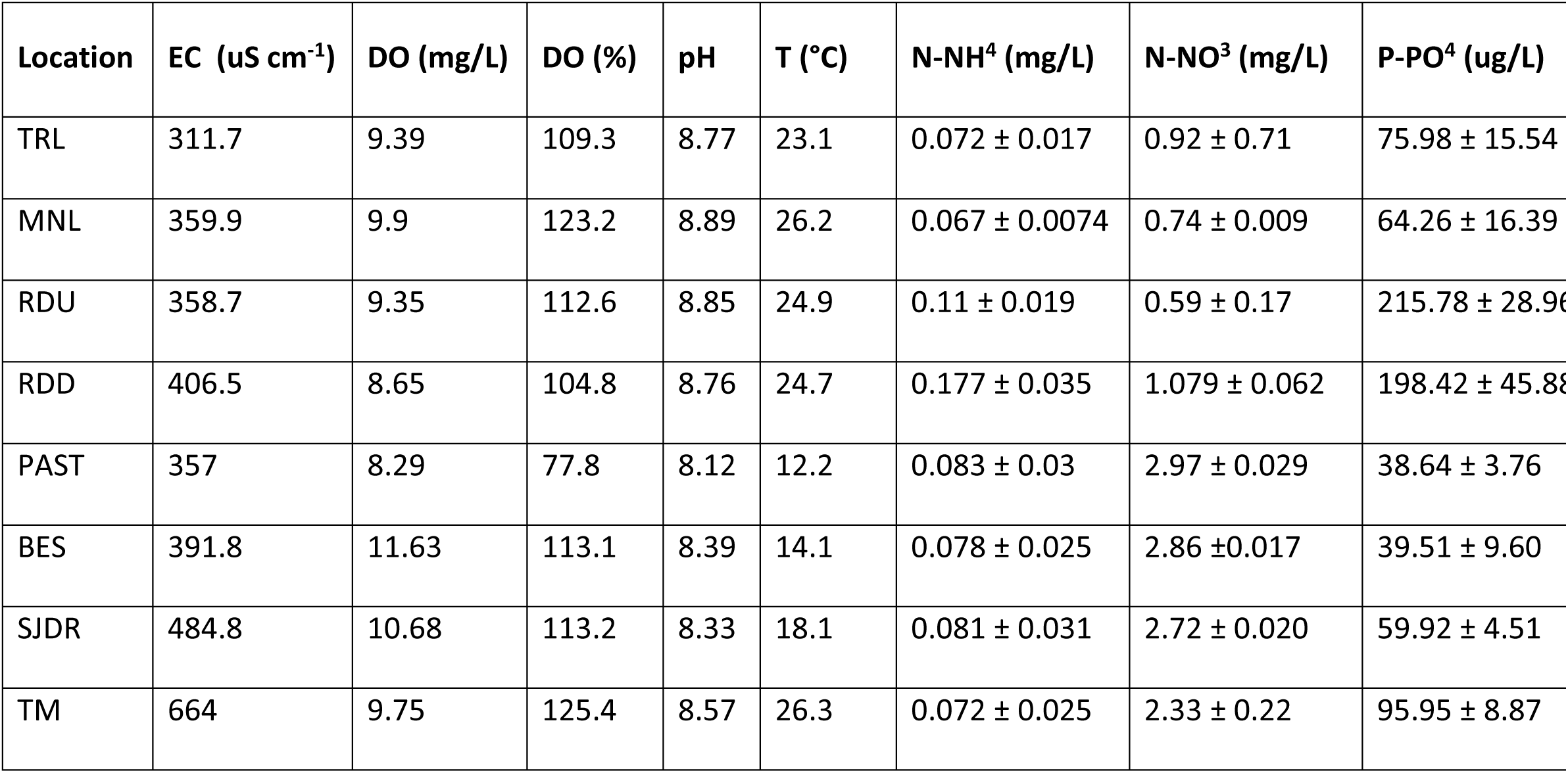
Physicochemical and nutrient parameters by location. EC = specific electrical conductivity (µS cm⁻¹ at 25 °C); DO = dissolved oxygen (mg L⁻¹) and O₂ saturation (%); T = temperature (°C). Nutrient concentrations (NH₄⁺-N, NO₃⁻-N in mg L⁻¹; PO₄³⁻-P in µg L⁻¹) are reported as mean ± SD

| Location | EC ($\mu\text{S cm}^{-1}$) | DO (mg/L) | DO (%) | pH | T ($^{\circ}\text{C}$) | N-NH<sup>4</sup> (mg/L) | N-NO<sup>3</sup> (mg/L) | P-PO<sup>4</sup> ($\mu\text{g/L}$) |
| --- | --- | --- | --- | --- | --- | --- | --- | --- |
| TRL | 311.7 | 9.39 | 109.3 | 8.77 | 23.1 | 0.072 ± 0.017 | 0.92 ± 0.71 | 75.98 ± 15.54 |
| MNL | 359.9 | 9.9 | 123.2 | 8.89 | 26.2 | 0.067 ± 0.0074 | 0.74 ± 0.009 | 64.26 ± 16.39 |
| RDU | 358.7 | 9.35 | 112.6 | 8.85 | 24.9 | 0.11 ± 0.019 | 0.59 ± 0.17 | 215.78 ± 28.96 |
| RDD | 406.5 | 8.65 | 104.8 | 8.76 | 24.7 | 0.177 ± 0.035 | 1.079 ± 0.062 | 198.42 ± 45.88 |
| PAST | 357 | 8.29 | 77.8 | 8.12 | 12.2 | 0.083 ± 0.03 | 2.97 ± 0.029 | 38.64 ± 3.76 |
| BES | 391.8 | 11.63 | 113.1 | 8.39 | 14.1 | 0.078 ± 0.025 | 2.86 ± 0.017 | 39.51 ± 9.60 |
| SJDR | 484.8 | 10.68 | 113.2 | 8.33 | 18.1 | 0.081 ± 0.031 | 2.72 ± 0.020 | 59.92 ± 4.51 |
| TM | 664 | 9.75 | 125.4 | 8.57 | 26.3 | 0.072 ± 0.025 | 2.33 ± 0.22 | 95.95 ± 8.87 |

### 3.3. Sequence data, rarefaction curves and high-level taxonomy overview

To ensure sufficient DNA extraction from all samples, experimental duplicates were conducted at the 8 sampling sites and later combined into the same pool after DNA extraction. After sequencing and filtering the samples, the 16S rRNA sequences obtained by shotgun metagenomics were retrieved, analysed, and taxonomically assigned. Given that the two datasets differ in target, sequencing technology, and analysis pipelines (full-length 16S amplicons on MinION vs. whole-genome shotgun on Illumina), cross-method results here are not intended as a head-to-head benchmark; we therefore avoid causal interpretations of differences between modalities and use the shotgun dataset to contextualize the field workflow.

For MinION sequencing, an average of 100,000 reads were obtained across all samples, with 92 ± 1.5 % identified as bacteria. In contrast, shotgun metagenomics approach yielded an average of 8,328,879 reads, which were initially split. However, after filtering the reads and associating them with the 16S rRNA region, only 0.05 ± 0.008 % of the sequences were taxonomically classified as bacteria (**Table S3**).

Species-level alpha-rarefaction clearly separates the behavior of both datasets (**Figure 3**). MinION full-length 16S amplicons show trajectories that approach an asymptote for most samples indicating that additional sequencing would yield diminishing returns in newly detected species. In contrast, shotgun metagenomes never approach a plateau in any sample within the obtained depths: richness continues to increase monotonically. These patterns are consistent with the fundamental differences between targeted 16S amplicons and untargeted whole-genome data and are not interpreted as superiority of one modality over the other.

**Figure 3.**
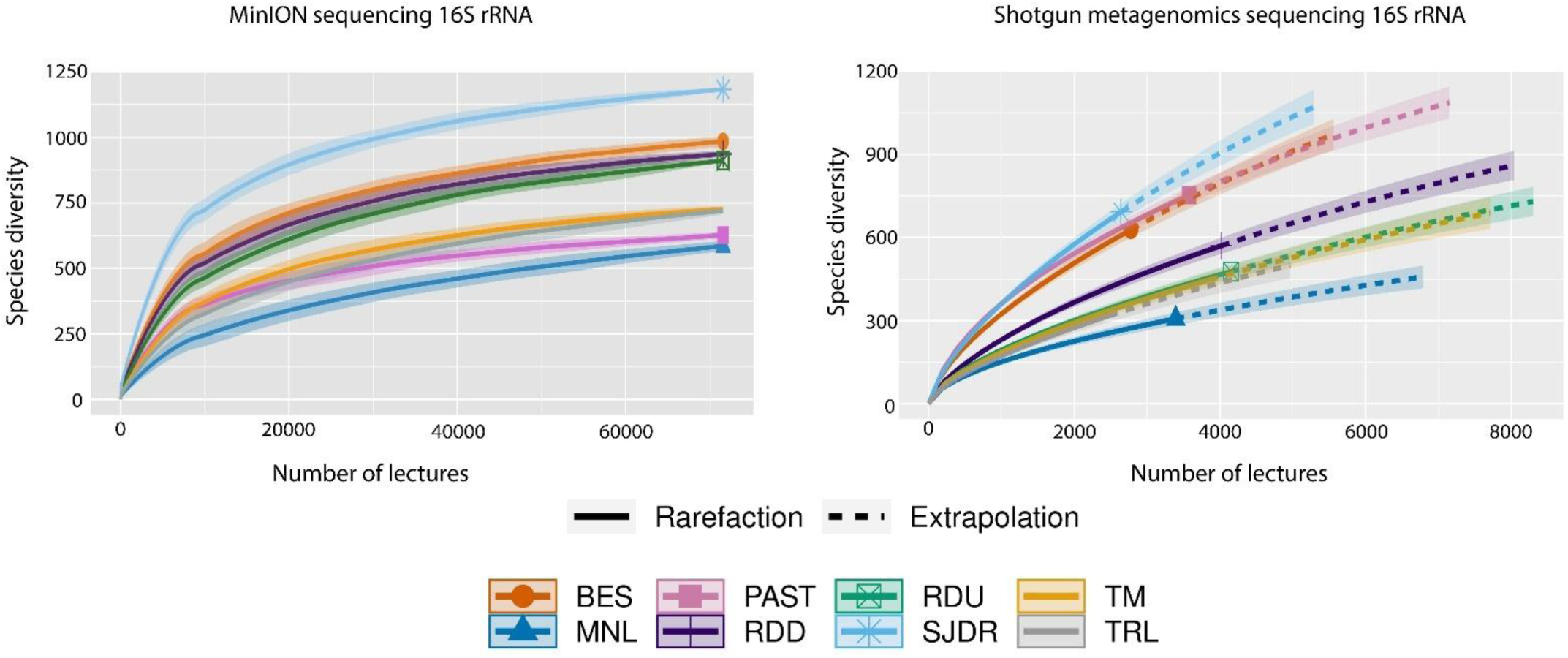
Species-level alpha rarefaction for MinION full-length 16S amplicons and Illumina shotgun metagenomes. Lines represent per-sample trajectories summarized as the median across 10 iterations at each of 40 depth steps; shaded ribbons indicate the interquartile range. Because the number of informative shotgun reads was low, we include a short extrapolated segment (dashed lines) to illustrate the predicted trend beyond the maximum observed depth.

Following taxonomic classification, bar graphs were plotted to visualize the most abundant taxonomic classes, highlighting differences between sequencing methodologies (**Figure 4**, **Table S4**). For MinION sequencing, the most abundant classes, regardless of sampling location, were Gammaproteobacteria (49.4 ± 15.6 %), Alphaproteobacteria (12.2 ± 3.6 %), and Bacilli (8.5 ± 19.6 %). Results also revealed clear distinctions between sampling locations. For example, at the PAST site, Bacilli was the major class (56.9 %), represented by the genus Exiguobacterium (55.8 %). In contrast, at TM site, the abundance of Actinobacteria was notably higher (19.9 %) compared to upstream sites. For shotgun metagenomics, results reflected substantially different outcomes. The most abundant classes identified were Betaproteobacteria (44.9 ± 12.3 %), Alphaproteobacteria (17.7 ± 5.5 %), and Actinobacteria (9.2 ± 3.5 %). At the PAST site, the relative abundance of Bacilli was reduced to 10.7 %, while at the TM site, the abundance of Actinobacteria (12.5 %) remainded higher than in the rest of the sampling sites.

**Figure 4.**
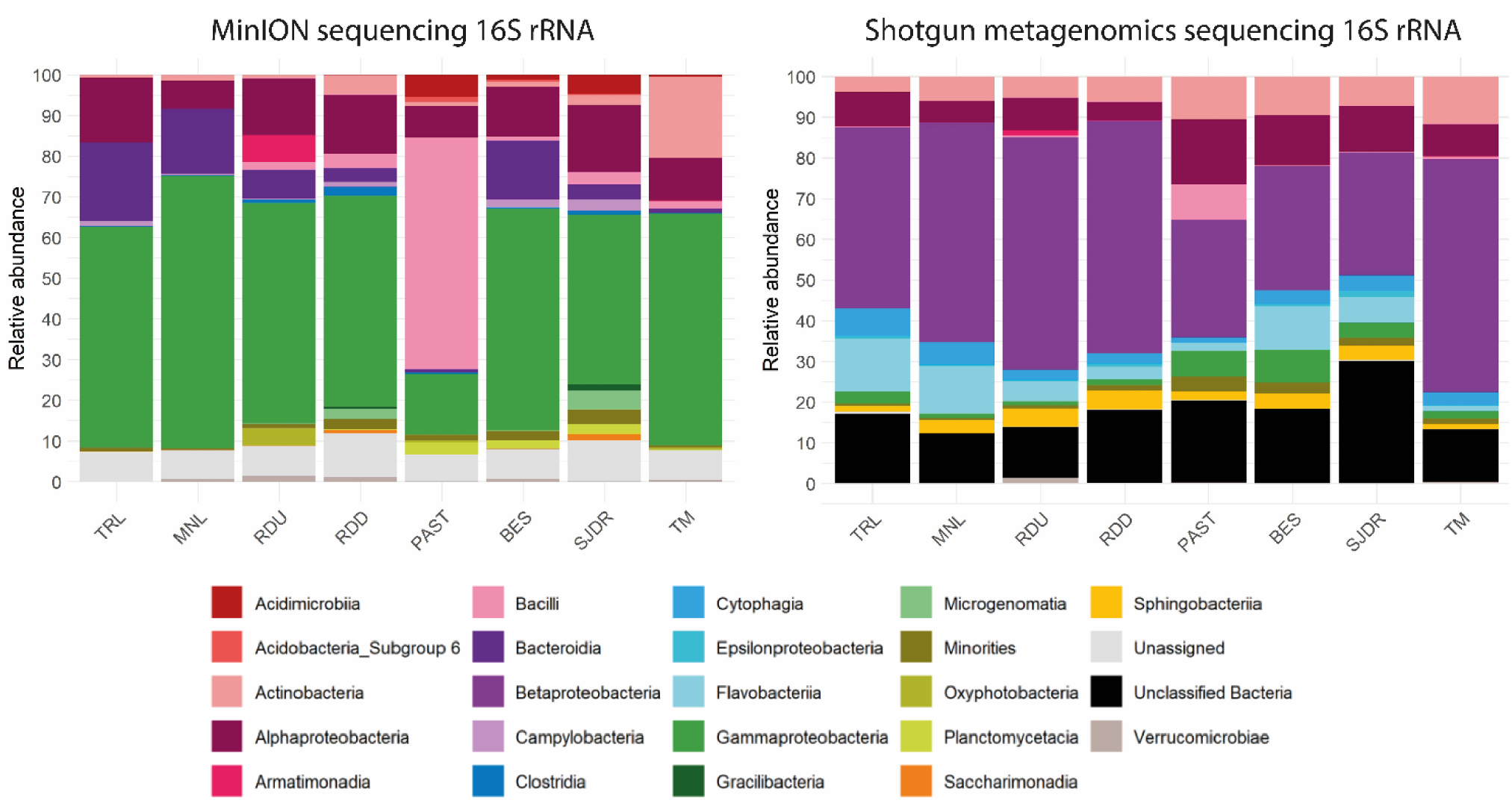
The graph on the left reflects the results obtained by MinION sequencing and the graph on the right reflects the results obtained by ILLUMINA Shotgun metagenomics. Both share the same legend and the minority species are those with a relative abundance of less than 1%.

### 3.4 Alpha and Beta-Diversity

Analyses of alpha and beta diversity were performed at genus level after taxonomic classification. Alpha diversity was assessed by calculating richness (R) and the Shannon-Wienner index (**Figure 5a**; **Table S5**).

**Figure 5.**
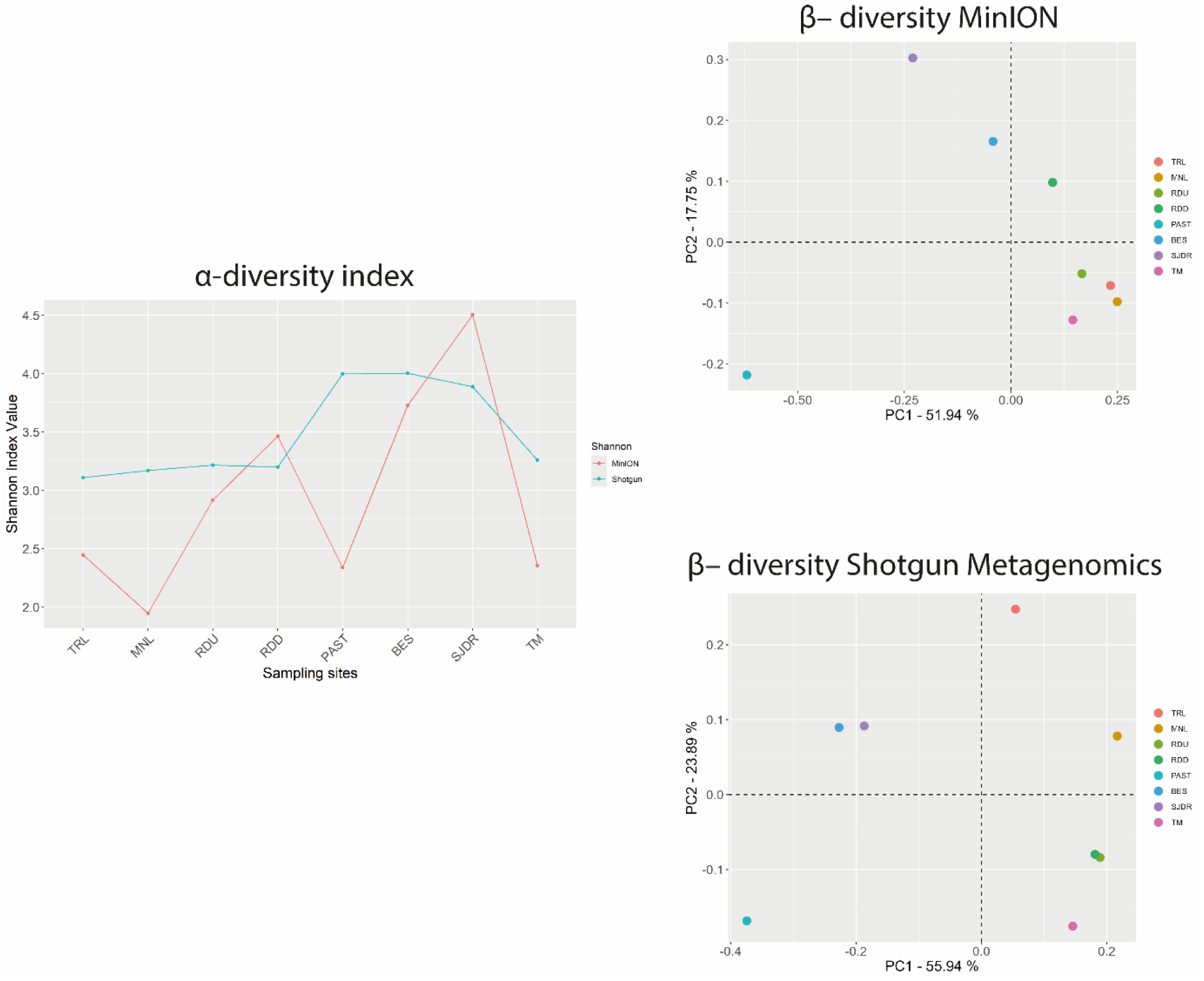
Alpha and beta diversity measures of genera detected using MinION and shotgun metagenomics. (a) Box plot of observed richness between sampling types. (b–d) Non-metric multidimensional scaling (NMDS) plots displaying ASV community structure between sample types and among sites for (b) 16S, (c) 18S and (d) COI genes.

The richness index was significantly higher for the 16S rRNA amplification performed by MinION (773.9 ± 204.3) than in shotgun metagenomics. In some locations, genome detection through amplification was up to 6 times higher, as observed at the SJR site (1099 by MinION amplicon vs 178 detected by shotgun metagenomics). Conversely, Shannon Wienner indicated lower diversity values for MinION sequenced samples (2.96 ± 0.87) compared to overall shotgun metagenomics results (3.48 ± 0.40). Despite this, MinION results showed greater dispersion, allowing more detailed characterization of specific diversity at each sampling site.

The Beta diversity results revealed that both sequencing methodologies showed a similar dispersion along the first axis (PC1), which explained the greatest differentiation between communities established at each site (51.94% for MinION sequencing and 55.94% for ILLUMINA Shotgun metagenomics (**Figures 5b, 5c**). Prokaryote communities belonging to the upstream sites (TRL, MNL, RDU, RDD) were primarily clustered on one side of the PC1 axis, while PAST, BES and SJDR downstream samples (i.e. excluding TM) were clustered at the opposite end. The second axis (PC2) accounted for significantly less variance (17.15% for MinION sequencing and 23.89% for shotgun metagenomics) and showed a dispersion pattern that varies between sites. For shotgun metagenomics, the close proximity of the RDU and RDD sampling sites was reflected in their minimal dispersion along PC2, consistent with the geographic proximity of these sites.

## 4. Discussion

The most recent studies on microbial diversity in rivers highlight the need to develop alternative and standardized methods for the rapid and efficient application of molecular techniques, allowing long-term monitoring of water bodies in Europe (Pascher et al., 2022). The results presented in this study offer an innovative solution, yielding viable results in less than 48 h and allowing the simple monitoring of a large number of taxa. Given that the MinION 16S amplicon dataset and the Illumina shotgun dataset differ in target, sequencing technology, and analytical pipelines, we present them side-by-side to contextualize the field workflow rather than to benchmark platforms; cross-method differences are not interpreted as platform performance

First, one of the main advantages ofthis study is the use of a USV for water quality monitoring and integrated samples collection. USVs are becoming an essential tool for water monitoring, although changes in regulations and the growing interest in improving monitoring strategies necessitate their continuous updates to meet current demands (Zhang et al., 2023). We have already shown the efficiency of USVs by taking measurements of different water quality parameters. Castellini et al. (2020) used a version of the USV to take continuous measurements of temperature, conductivity, altitude and dissolved oxygen with a multivariable sensor in different lakes and rivers in Spain and Italy during the 5.4 h of battery life. Katsouras et al. (2024) showed the economic feasibility of implementing this model in Lake Koumoundourou and the Acheloos, Asopos and Kifissos rivers of Greece to measure spatial fluctuations of α-chlorophyll, electrical conductivity, dissolved oxygen and pH.

In this study, the INCATCH USV model had been upgraded including an advanced filtration system which has provided a great advantage not available in previous existing models.

Powers et al. (2018) focused on the development of USVs capable of collecting water samples at different depths in a lake (5 and 50 cm) and then providing average bacterial concentrations (CFU/ml) for KBC and TSA medium. Preston et al. (2024) developed a USV that was able to collect 52 samples along a 4200 km transect, which were then used to study the associated eukaryotic communities by amplification. However, they found that the samples were not immediately extracted and properly preserved, resulting in loss of relevant DNA concentration and limiting the technology.

However, the results presented in this study prove that the protocol developed is highly efficient in several ways. Firstly, the sequential arrangement of filters with different pore sizes, allows at the same time to separate the particulate fraction and only collect the free-living planktonic (Amalfitano et al., 2017) microorganisms avoiding the double filtration commonly applied in the laboratory (Amalfitano et al., 2017) and preventing potential cross contamination during transport and samples manipulation.

Secondly, the filtering capacity of the USV avoids the transfer of large volumes of water for subsequent filtering in the lab, which can involve up to 6 (or more) liters of water (Hunter et al., 2019) depending on trophic conditions of the ecosystems, and allow collecting greater amount of samples from different sites of the same transect, ensuring the representativeness of the variability that may occur along longer river stretches..

The second advantage of implementing the protocols used within this study is the development of a portable lab, which significantly decreases the time needed for the analysis. The feasibility of implementing a portable laboratory has already been tested in previous studies (Acharya et al., 2019; Hu et al., 2018), with promising results. Acharya et al. (2020) evaluated by 16S MinION amplicon of treated urban wastewater from a plant located in the UK and from an Ethiopian river, obtaining reliable results on microbial communities in only 3 days. Hu and colleagues (2018) used MinION 16S rRNA sequencing compared to shotgun and to *E. coli* culture to detect faecal bacteria in sewage systems, showing a clear correlation between the three technologies, although with notable differences in the amount of information provided by each.

These studies indicate that MinION is currently the most widely deployed long-read platform for fully portable field workflows. In our case, we have reduced the time from sampling to data analysis to less than 2 days. Furthermore, regardless of whether shotgun or amplicons analyses are performed, MinION studies significantly reduce the time and the probability of cross contamination during transport or in the lab by performing the sample processing and analyses in situ just after collection (Acharya et al., 2020; Hu et al., 2018). In cost terms, our portable ONT workflow (≈900 € per sample for single-sample runs; ≈ 93 € per sample when multiplexing 12) sits between targeted qPCR and high-throughput laboratory sequencing. Peer-reviewed studies show that MinION enables field-deployable workflows with same-day to ∼24 h turnaround (Urban et al., 2021) whereas centralized Illumina pipelines typically minimize per-sample costs when large batches are pooled (van der Reis et al., 2023). Public core-facility rate cards illustrate typical ranges: 16S (Illumina) sequencing ∼ 100–110 US$ per sample, with library prep often billed separately ∼ 53–94 US$ per sample (Emory Integrated Genomics Core, 2025; Northwestern University Center for Genetic Medicine NUSeq Core, 2025; Weill Cornell Microbiome Core, 2025); shotgun metagenomics commonly ∼285–500 US$ per sample at shallow–moderate depths (MD Anderson Microbiome Core Facility, 2025). In our field consumables, the flow cell is the dominant driver of cost, explaining the strong economies of scale with multiplexing. Practically, when dozens of samples can be pooled and shipped to a core, core-based 16S often minimizes per-sample consumables; conversely, for incident response, remote sites, or limited batching, the field-deployable ONT approach is cost-reasonable while uniquely enabling ∼12–24 h on-site time-to-result.

However, this approach also presents some problems and limitations, such as the need to keep part of the materials at 4°C, specifically PCR reagents and MinION flow cells, which can represent a barrier to application in extremely warm conditions or can limit the time allowed for sample collection decreasing the possibility to integrate long stretches over long periods. Furthermore, in the application of MinION nanopore sequencing, the most relevant issue was related to the error rate which was significantly higher than other sequencing techniques (i.e. Illumina). However, this limitation seems to have been overcome in recent years in which the accuracy MinION sequencing has increased from 60% to 90% (Rang et al., 2018). Because our data were generated on R9.4.1 flow cells, our use of Guppy (v4.0.11) at the time of analysis aligned with ONT’s model support for this chemistry. To reduce error-driven inflation of apparent diversity, we applied stringent filters (PHRED ≥12; full-length 16S window 1,400–1,500 bp), performed chimera removal, and summarized composition at the genus level using the MetONTIIME workflow. We also contrasted these profiles with Illumina shotgun metagenomes as an orthogonal check of community patterns. As a sensitivity analysis, future work could re-basecall raw signals with the last Dorado release supporting R9.4.1 to confirm robustness of the main ecological conclusions.

Because approach (amplicon vs. shotgun), platform (ONT vs. Illumina), and pipelines differ by design, we interpret cross-dataset contrasts cautiously and use the shotgun dataset as a complementary laboratory reference. Our goal is to assess whether the on-site workflow recovers the same environmental patterns (e.g., reach-scale gradients) under field conditions, rather than to perform a head-to-head platform benchmark.

On other hands, this portable sequencing instrument present also notable advantages, for example it allows the sequencing of remarkably long sequences that are significantly longer than ILLUMINA light (Werner et al., 2022) This factor should be considered when carrying out simple monitoring or searching for much more deep and precise information in environmental samples.

In comparison with previous field-based MinION applications, our work presents several distinctive advances. (Maestri et al., 2019) demonstrated the feasibility of a portable MinION workflow for biodiversity tracking, while (Johnson et al., (2017) successfully applied nanopore sequencing in extreme polar environments to characterize microbial communities under challenging logistical conditions. More recently, (Reddington et al., (2020) applied MinION sequencing to freshwater monitoring across multiple rivers, highlighting issues related to large water volumes and logistical constraints, and (Werner et al., 2022b) emphasized that fully portable workflows for aquatic research remain rare, particularly in terms of integrating all steps from biomass collection to sequencing. While these studies proved the adaptability of MinION in the field, they were constrained by sample handling requirements, incomplete portability, or preservation challenges. In contrast, our study integrates an upgraded USV equipped with sequential in situ filtration, immediate DNA processing, and portable MinION sequencing. This approach minimizes DNA degradation, reduces handling steps, and enables standardized long-term monitoring across extended river transects, representing a clear methodological advance beyond existing field-based MinION efforts.

Short-term and reach-scale variability in environmental conditions can shape riverine microbiomes and therefore confound sequencing-based contrasts across locations. In our study, we found an important different gradient between the sampling sites at the level of conductivity, temperature, dissolved oxygen and nutrients. Such gradients are known to restructure bacterioplankton via environmental selection and hydrological control (Niño-García et al., 2016). For this reason, hydrological properties are used to predict the composition of functional microbial groups (Clark et al., 2022) and the recurring importance of temperature, pH, dissolved oxygen and phosphate as correlates of community turnover in impacted systems (Jeffries et al., 2016). Likewise, nitrogen and phosphorus enrichment underpin eutrophication responses with consequences for algal–microbial dynamics and oxygen availability (Dodds & Smith, 2016). Our design constrained methodological variability by integrating water along a 100 m transect at fixed 10 cm depth, standardizing filtration volume/time, pooling duplicate filters to stabilize DNA yield, processing all samples with a single laboratory/bioinformatic workflow with equal-depth summaries, and logging physicochemical covariates at each location. While these measures reduce bias, the absence of independent per-location biological replicates leads us to emphasize descriptive contrasts rather than formal hypothesis tests; future campaigns will include replicated time points and within-location replicates and apply multivariate models to partition environmental from methodological variance more explicitly. Under our sequencing depths and analysis settings, full-length 16S amplicons detected more taxa, which is expected for a targeted marker-gene assay; we do not interpret this as evidence of platform superiority. Shotgun metagenomics provided broader functional context (e.g., gene content, metabolic potential, resistance/virulence markers), complementing the field workflow. In this way, shotgun metagenomics provides much more information on the functionality of the microbial community, whereas the use of amplicons will be much more accurate for assessing changes in community structure and diversity and to detect and identify possible drivers of microbiological contamination in freshwater ecosystems. Furthermore, the species-level alpha rarefaction obtained revealed that full-length 16S MinION curves tended toward asymptotes for most samples—indicating diminishing returns within our depth range—whereas shotgun curves did not plateau, consistent with the dilution of informative reads outside the 16S rRNA locus. Summarizing at the species level (rather than ASVs) harmonizes resolution across technologies and reduces richness inflation from intragenomic variation and residual errors, improving ecological interpretability for management applications (Gotelli & Colwell, 2001; Tessler et al., 2017).

Although MinION has become increasingly reliable, it is important to acknowledge platform-specific limitations and trade-offs. Nanopore reads remain prone to insertion/deletion errors in homopolymeric regions, which can inflate apparent richness and complicate species-level resolution (Delahaye & Nicolas, 2021; Rang et al., 2018; Tyler et al., 2018), whereas Illumina shotgun sequencing yields lower per-base error rates but requires longer turnaround times and centralized facilities. Our results are consistent with these contrasts: genus-level profiles showed high concordance between platforms, but fine-scale assignments diverged. Such differences emphasize that MinION excels in rapid assessments of community structure, while shotgun metagenomics provides greater resolution of functional potential. Similar field-based studies have reported analogous trade-offs between speed and accuracy (Reddington et al., 2020; Urban et al., 2021). In practical terms, the rapid response capability we demonstrate—moving from USV-based sampling to taxonomic profiles in less than two days—can be directly linked to management scenarios such as early detection of upstream sewage inputs, tracing fecal contamination sources, or identifying incipient cyanobacterial blooms. These applications illustrate how portable sequencing can complement existing monitoring frameworks by providing actionable information on time scales relevant for environmental decision-making.In addition to the advantages of implementing MinION sequency technology, there is also the possibility to apply the bioinformatics protocol in a single step, through the development of the MetONTIIMEMetONTIIME workflow (Matoute et al., 2024), a protocol that, although more limited than the most commonly used bioinformatics processes such as Mothur (Schloss et al., 2009) or QIIME 2 (Bolyen et al., 2019), allows users without extensive knowledge, further streamlining the process and obtain clear results in a reduced time, overall facilitating its reproducibility. Eventually, the possibility of uploading these data to common databases, such as the National Center for Biotechnology Information platform, would allow new developments in databases to be applicable in the future with these data, as well as the results to be comparable in the future if new research is conducted in the same areas.

## 5. Conclusions

A mobile genomics laboratory has been set up and validated to perform cost effective microbiological monitoring of water samples in different field conditions and scenarios. The USV allows for quick and easy sampling of a large surface area, making possible to concentrate the sampling on specific sampling sites of interest or, on the contrary, to have an integrated sample representative of the whole selected area over a certain period of time. The procedure for using the genomic tool has been completely optimized and simplified so also stakeholders and citizen scientists could utilize it after appropriate training. Moreover, the inclusion of a simple bioinformatics protocol, applicable in a single step and with results easily analyzable through interactive graphs, makes it readily available to scientific community, water managers, public authorities responsible for catchment management, and the general public. Nevertheless, the method proposed has some limitations by resorting to MinION sequencing (lower sequence quality with a higher error rate but a very long sequence reading potential). Our field-deployable workflow recovered ecologically coherent patterns across river sections, while laboratory shotgun metagenomics provided complementary functional context. Rather than a head-to-head comparison, we illustrate how both modalities can be combined: MinION-16S for rapid, on-site community profiling in the field, and Illumina shotgun for in-depth functional characterization when centralized facilities are available. This complementary strategy supports long-term monitoring by reducing delays and enabling standardized, spatially resolved surveys without reliance on centralized infrastructure.

## Authors’ contributions

All authors have contributed intellectually and to the writing of this manuscript. All authors read and approved the final manuscript.

## Acknowledgements

MS was supported by a Juan de la Cierva-Formación contract (FJC2021-046825-I) from the Spanish Ministry of Science and Innovation. SMC was supported by a Investigo Programme contract funded by the European Union, Next Generation EU (grant number 2022 INV-1 00037).

