## Supplementary data for "On-site microbial community analysis in rivers: integrating autonomous sampling with a portable sequencing workflow"

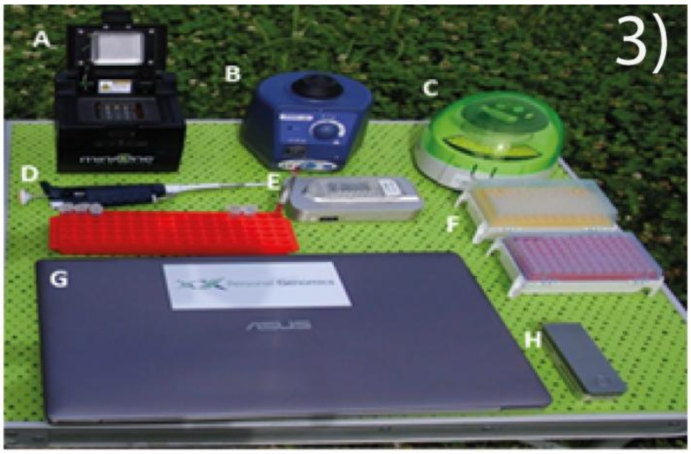


**Figure S1.** Close-up of lab components: A) thermal cycler; B) vortex mixer; C) microcentrifuge; D) micropipettes; E) fluorometer; F) filter tips; G) laptop; H) MinION™ sequencer.

| Activity Item | Item | Company |
|---|---|---|
| Portable Lab | Notebook with analysis software | ASUS |
| | MinION sequencer | Oxford Nanopore |
| | Pipet P20 | Gilson |
| | Pipet P200 | Gilson |
| | Pipet P2000 | Gilson |
| | Vortex | Scientific industries |
| | QuBit spectrophotometer | ThermoFisher |
| | Mini-centrifuge | Labnet Prism |
| | MinION portable PCR | MinION |

**Table S1.** List of equipment included in the portable laboratory.

| Activity | Consumable item | Company |
|---|---|---|
| DNA extraction<br>DNA amplification & sequencing | QuBit DsDNA BR Assay Kit | ThermoFisher |
| | DNeasy PowerWater Kit | Qiagen |
| | 16S Rapid Amplicon Barcoding Kit | Oxford Nanopore |
| | Agencourt AMPure XP | Beckman New England |
| DNA extraction | LongAmp Taq | Biolabs |
| | SpotON Flow Cell R9.4 | Oxford Nanopore |

**Table S2.** Consumables needed to carry out each of the tasks.

| **MinION** | | | | | | | | |
|---|---|---|---|---|---|---|---|---|
| Sequences | TRL | MNL | RDU | RDD | PAST | BES | SJDR | TM |
| Total reads | 1000000 | 1000000 | 1000000 | 1000000 | 1000000 | 1000000 | 1000000 | 1000000 |
| Reads associated with 16S rRNA | 73240 | 73320 | 101350 | 74380 | 108310 | 66250 | 70520 | 72400 |
| Non Bacteria Reads | 926760 | 926680 | 898650 | 925620 | 891690 | 933750 | 929480 | 927600 |
| Unclassified Bacteria reads | 10 | 10 | 0 | 0 | 0 | 0 | 0 | 0 |
| **ILLUMINA Shotgun Metagenomics** | | | | | | | | |
| Reads | TRL | TM | SJR | RDU | RDD | PAST | MNL | BES |
| Total reads | 7436298 | 9087645 | 8365547 | 9045371 | 8445418 | 8985181 | 7415341 | 7850236 |
| Reads associated with 16S rRNA | 3062 | 4852 | 3365 | 5183 | 5161 | 4436 | 4239 | 3417 |
| Non Bacteria Reads | 18 | 6 | 15 | 6 | 14 | 2 | 1 | 3 |
| Unclassified Bacteria reads | 3044 | 4846 | 3350 | 5177 | 5147 | 4434 | 4238 | 3414 |

**Table S3.** Summary of the results obtained after sequencing, filtering and taxonomic classification by ILLUMINA Shotgun and MinION amplicon sequencing**.**

| MinION | | | | | | | | |
|---|---|---|---|---|---|---|---|---|
| Classes | TRL | MNL | RDU | RDD | PAST | BES | SJDR | TM |
| Gammaproteobacteria | 54,09 | 66,96 | 54,14 | 51,69 | 14,83 | 54,48 | 41,70 | 56,91 |
| Alphaproteobacteria | 15,81 | 6,85 | 13,93 | 14,55 | 7,67 | 12,21 | 16,43 | 10,45 |
| Bacilli | 0,04 | 0,03 | 1,91 | 3,41 | 56,88 | 0,97 | 2,88 | 1,84 |
| Bacteroidia | 19,33 | 15,92 | 6,99 | 3,47 | 0,86 | 14,42 | 3,80 | 1,07 |
| Unassigned | 7,32 | 7,05 | 7,44 | 10,83 | 6,63 | 7,24 | 10,14 | 7,33 |
| Actinobacteria | 0,75 | 1,48 | 0,87 | 4,73 | 1,10 | 1,18 | 2,53 | 19,87 |
| Acidimicrobiia | 0,01 | 0,01 | 0,03 | 0,06 | 5,44 | 1,26 | 4,63 | 0,48 |
| Planctomycetacia | 0,04 | 0,01 | 0,16 | 0,16 | 3,02 | 1,97 | 2,39 | 0,21 |
| Campylobacteria | 1,26 | 0,37 | 0,35 | 1,18 | 0,02 | 2,01 | 2,73 | 0,01 |
| Microgenomatia | 0,07 | 0,03 | 0,10 | 2,60 | 0,03 | 0,05 | 4,58 | 0,04 |

| | | | | | | | | |
|---|---|---|---|---|---|---|---|---|
| Armatimonadia | 0,01 | 0,01 | 6,58 | 0,04 | 0,01 | 0,01 | 0,00 | 0,16 |
| Clostridia | 0,19 | 0,14 | 0,80 | 2,25 | 0,40 | 0,37 | 1,03 | 0,19 |
| Oxyphotobacteria | 0,04 | 0,01 | 4,08 | 0,06 | 0,49 | 0,04 | 0,01 | 0,35 |
| Verrucomicrobiae | 0,05 | 0,72 | 1,42 | 1,13 | 0,07 | 0,80 | 0,08 | 0,41 |
| Saccharimonadia | 0,03 | 0,03 | 0,04 | 0,76 | 0,02 | 0,08 | 1,52 | 0,03 |
| Gracilibacteria | 0,05 | 0,01 | 0,08 | 0,59 | 0,02 | 0,07 | 1,53 | 0,03 |
| Acidobacteria_Subgroup 6 | 0,02 | 0,01 | 0,02 | 0,07 | 1,16 | 0,49 | 0,40 | 0,05 |
| Minorities | 0,89 | 0,36 | 1,08 | 2,44 | 1,36 | 2,36 | 3,62 | 0,59 |
| Shotgun metagenomics | | | | | | | | |
| Classes | TRL | MNL | RDU | RDD | PAST | BES | SJDR | TM |
| Betaproteobacteria | 44,48 | 53,86 | 57,23 | 57,06 | 28,94 | 30,55 | 30,19 | 57,23 |
| Unclassified Bacteria | 17,08 | 12,24 | 12,54 | 17,90 | 20,24 | 18,29 | 29,99 | 12,82 |
| Alphaproteobacteria | 8,43 | 5,38 | 8,03 | 4,73 | 15,94 | 12,29 | 11,14 | 7,91 |
| Actinobacteria | 3,89 | 6,02 | 5,19 | 6,12 | 10,57 | 9,54 | 7,25 | 11,64 |
| Flavobacteriia | 13,00 | 11,75 | 4,94 | 3,18 | 1,94 | 10,65 | 6,39 | 1,30 |
| Cytophagia | 6,79 | 5,57 | 2,49 | 2,60 | 1,31 | 3,31 | 3,92 | 3,24 |
| Gammaproteobacteria | 2,97 | 1,09 | 1,00 | 1,38 | 6,38 | 7,99 | 3,68 | 1,85 |
| Sphingobacteriia | 1,31 | 3,28 | 4,50 | 4,59 | 2,12 | 3,78 | 3,48 | 1,36 |
| Bacilli | 0,13 | 0,00 | 0,39 | 0,04 | 8,63 | 0,18 | 0,27 | 0,66 |
| Epsilonproteobacteria | 0,62 | 0,26 | 0,25 | 0,66 | 0,05 | 0,64 | 1,34 | 0,10 |
| Verrucomicrobiae | 0,00 | 0,02 | 1,29 | 0,10 | 0,23 | 0,00 | 0,00 | 0,43 |
| Unassigned | 0,59 | 0,02 | 0,12 | 0,27 | 0,05 | 0,09 | 0,45 | 0,12 |
| Armatimonadia | 0,03 | 0,00 | 1,25 | 0,04 | 0,00 | 0,00 | 0,00 | 0,10 |
| Minorities | 0,69 | 0,52 | 0,79 | 1,34 | 3,61 | 2,69 | 1,90 | 1,22 |

**Table S4.** The abundance of the most abundance classes using Shotgun metagenomics and MinION amplicon sequencing in all locations.

| Shannon | TRL | MNL | RDU | RDD | PAST | BES | SJDR | TM |
|---|---|---|---|---|---|---|---|---|
| MinION | 2,44 | 1,94 | 2,92 | 3,46 | 2,33 | 3,73 | 4,50 | 2,35 |
| Shotgun | 3,11 | 3,17 | 3,21 | 3,20 | 4,00 | 4,00 | 3,89 | 3,26 |
| Richness | TRL | TM | SJR | RDU | RDD | PAST | MNL | BES |
| MinION | 717 | 681 | 1099 | 912 | 852 | 503 | 528 | 899 |
| Shotgun | 107 | 173 | 178 | 162 | 172 | 197 | 115 | 193 |

**Table S5.** Estimation of alpha diversity index using Shannon-Wienner Index and Richness (number of genus).

**Supplementary protocol S1.** MetONTIIME pipeline (16S rRNA minion profiling)

```
metontiime run \
        --input samplesheet.csv \
        --minQual 12 \
        --MinReadLength 1400 \
       --MaxReadLength 1500 \
        --maxNumReads 100000 \
```

```
--seed 42 \
--classifier vsearch \
--ref_fasta silva_132_SSURef_NR99.fasta \
--ref_taxonomy silva_132_taxonomy.txt \
--outdir metontiime_out
```
